# Reduced sphingolipid biosynthesis modulates proteostasis networks to enhance longevity

**DOI:** 10.1101/2022.05.20.492522

**Authors:** Nathaniel L. Hepowit, Eric Blalock, Sangderk Lee, Jason A. MacGurn, Robert C. Dickson

**Author notes:** Correspondence Robert C. Dickson, Department of Molecular and Cellular Biochemistry, University of Kentucky, 741 South Limestone, Lexington, KY, USA 40536., Jason A. MacGurn, Depart of Cell and Developmental Biology, Vanderbilt University, PMB 407935, 465 21st Avenue South, Medical Research Building III, Room 4130, Nashville, TN 37240-7935.

## Abstract

As the elderly population increases, chronic, age-associated diseases are challenging healthcare systems around the world. Nutrient limitation is well known to slow the aging process and improve health. Regrettably, practicing nutrient restriction to improve health is unachievable for most people. Alternatively, pharmacological strategies are being pursued including myriocin which increases lifespan in budding yeast. Myriocin impairs sphingolipid synthesis, resulting in lowered amino acid pools which promote entry into a quiescent, long-lived state. Here we present transcriptomic data during the first 6 hours of drug treatment that improves our mechanistic understanding of the cellular response to myriocin and reveals a new role for ubiquitin in longevity. Previously we found that the methionine transporter Mup1 traffics to the plasma membrane normally in myriocin-treated cells but is not active and undergoes endocytic clearance. We now show that *UBI4*, a gene encoding stressed-induced ubiquitin, is vital for myriocin-enhanced lifespan. Furthermore, we show that Mup1 fused to a deubiquitinase domain impairs myriocin-enhanced longevity. Broader effects of myriocin treatment on ubiquitination are indicated by our finding of a significant increase in K63-linked ubiquitin polymers following myriocin treatment. Although proteostasis is broadly accepted as a pillar of aging, our finding that ubiquitination of an amino acid transporter promotes longevity in myriocin-treated cells is novel. Addressing the role of ubiquitination/deubiquitination in longevity has the potential to reveal new strategies and targets for promoting healthy aging.

## 1 INTRODUCTION

Aging is widely accepted as a major risk factor for many chronic diseases and resultant physiological decline leading to mortality (Gonzalez-Freire et al., 2020). Research on many fronts is revealing potential ways to postpone age-related decline, maintain normal physiological function longer and improve healthspan. Some of the most promising research seeks to limit nutrient intake or increase daily fasting time as a means to improve healthspan in humans (Aon et al., 2020; Madeo, Carmona-Gutierrez, Hofer, & Kroemer, 2019; Stekovic et al., 2019). Still, these strategies will be difficult for most humans to adhere to in order to gain health benefits. Pharmacological agents offer a potential way to obtain the beneficial effects of nutrient limitation, but such compounds have yet to be identified although progress is encouraging (Gonzalez-Freire et al., 2020; Lee, Hill, Bitto, & Kaeberlein, 2021; Tyshkovskiy et al., 2019).

We have identified a potential pharmacological agent, the natural product myriocin (Myr, ISP-1), that increases chronological lifespan in budding yeasts (*Saccharomyces cerevisiae*) by more than two-fold (N. L. Hepowit et al., 2021). Myr works, at least in part, by reducing the free pool of most amino acids similar to what amino acid restriction does (Green & Lamming, 2019; Lee et al., 2021; Longo et al., 2015). Our interest in Myr stems from its target enzyme serine palmitoyltransferase (SPT), catalyzing the first and rate limiting step in sphingolipid biosynthesis in all eukaryotes (Dickson & Lester, 1999; Hanada, 2003; Miyake, Kozutsumi, Nakamura, Fujita, & Kawasaki, 1995). In addition, Myr was first identified in a search for antibiotics (Kluepfel, Bagli, Baker, Charest, & Kudelski, 1972) and anti-inflammatory drugs (Fujita et al., 1994). More recently, it has shown beneficial effects in treating age-associated diseases including diabetes, cancers, neurological and cardiovascular disorders (Lin, Wang, Marcogliese, & Bellen, 2019; Petit et al., 2020; Piano et al., 2020; Quinville, Deschenes, Ryckman, & Walia, 2021; Tippetts, Holland, & Summers, 2021; Woo et al., 2019) and other diseases including muscular dystrophies, cystic fibrosis and retinopathy (Laurila et al., 2022; Mingione et al., 2021; Shiwani et al., 2021)

Sphingolipids serve as both structural components of cellular membranes and as signal or regulatory molecules influencing many physiological processes, particularly in mammals (Dickson, 2008; Hannun & Obeid, 2008; Merrill, 2011; Quinville et al., 2021). Because most de novo lipid biosynthesis begins in the endoplasmic reticulum and continues in the Golgi Apparatus before the terminal products are distributed to cellular membranes, Myr treatment has the potential to diminish or enhance a variety of processes. Our recent studies identified diminished processes that may foster longer lifespan: newly synthesized Mup1, the major high-affinity methionine (Met) transporter, trafficked normally to the plasma membrane (PM) but was inactive in drug-treated cells resulting in reduced Met uptake as the fraction of active Mup1 was diluted by cell growth and division. Moreover, Myr-promoted endocytic clearance of Mup1 indicating that altering sphingolipid levels triggers remodeling of nutrient transporter composition at the PM (N. L. Hepowit et al., 2021). Thus, post-translational effects are vital to Myr-induced down-sizing of amino acid pools.

Previously we found that Myr treatment had large, global effects on transcription after 6-7 cell doublings (Liu et al., 2013). In the present work, we examined mRNA levels during the initial stages of Myr treatment to construct an overview of transcriptional changes with the aim of identifying how long it takes cells to respond to drug treatment and to determine if transcription plays a prominent role in lowering amino acid pools. Additionally, we sought to identify novel factors critical for Myr-enhanced lifespan. We find that very few transcripts respond to Myr in the first 4 h of treatment, but thereafter transcription is strongly up-regulated. We find no indication of transcription having a prominent role in maintaining low amino acid pools in Myr-treated cells. However, transcript data suggested a novel role for ubiquitin in lifespan and targeted studies identified ubiquitination of Mup1 as essential for Myr-enhanced longevity.

## 2 RESULTS

### 2.1 Four hours of myriocin treatment induce robust transcriptional changes

To examine transcriptional changes induced by Myr treatment, we diluted stationary phase cells (50-60% in G_o_ or quiescent, Q, phase) into fresh culture medium (Time 0), with and without Myr treatment. Samples for analysis of mRNA abundance by RNA seq were taken at 1 h intervals over a 6 h time course (Figure 1A). Data for the zero time were omitted from this analysis because of bias in library sizes which were smaller than expected if most genes were not differentially expressed. The normalized RNA seq data for the 1-6 h time course contained transcripts, expressed as transcripts per million (TPM), mapped to 6198 genes, 5169 of which were uniquely annotated and of sufficient signal intensity for subsequent analysis (File S1). Filtered data were analyzed by two-way ANOVA with a statistical cutoff of ρ = ≤ 0.01 to give a set of 4964 significant genes (Figure 1B). These were further sorted into drug (D), time (T), both drug and time (DxT) and interactions (TxDxI) (Figure 1C, Venn diagram).

**FIGURE 1.**
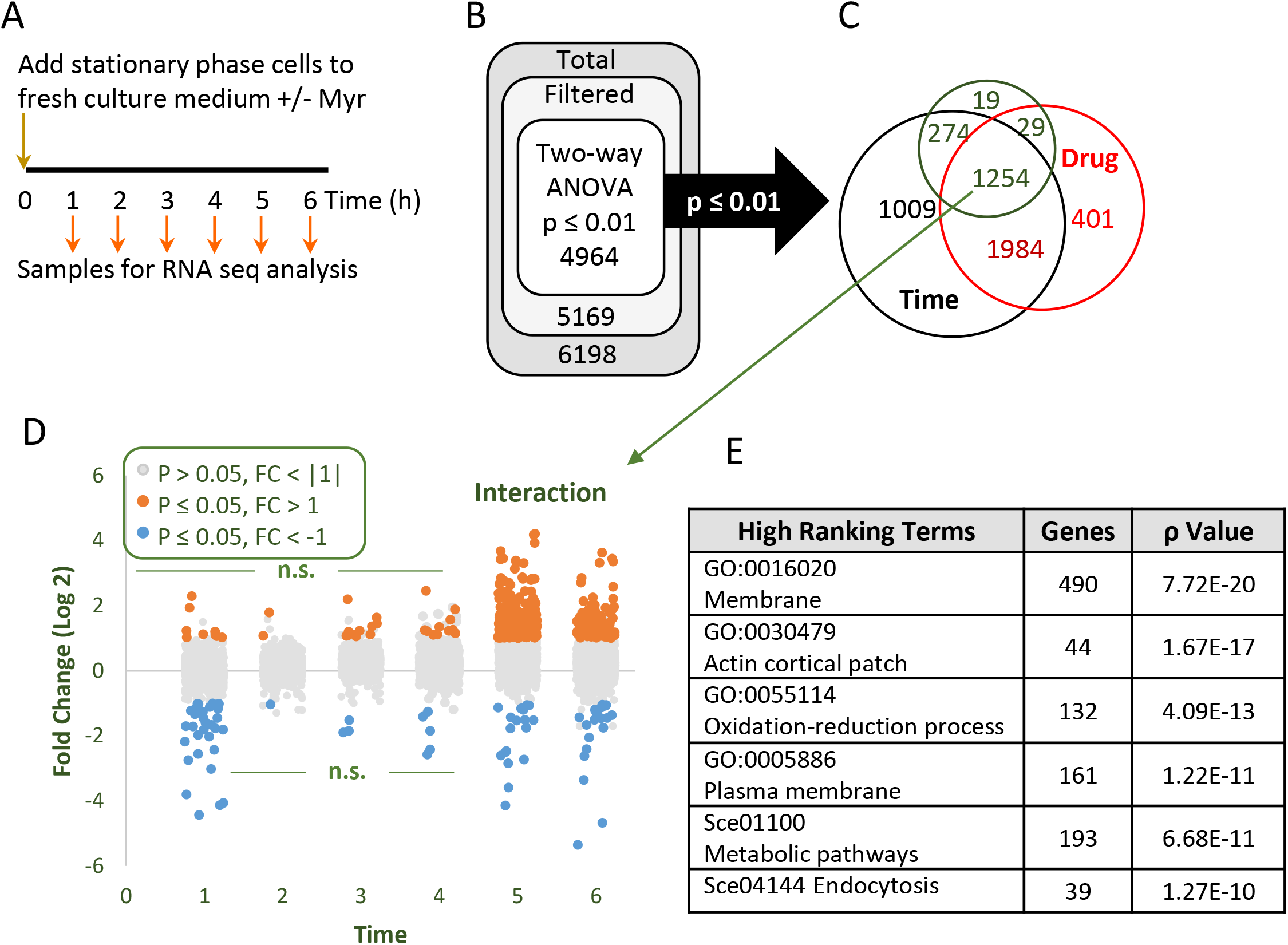
Summary of transcriptomics data. (a) Outline of experimental design for collecting cells, treated or not treated with Myr, for analysis of mRNAs by RNA seq. (b) Overview of RNA seq data analysis by two-way ANOVA. (c) Venn diagram summarizing the results of the ANOVA analysis for Drug, Time and Interaction effects. (d) Diagram summarizing analysis of the 1254 genes (transcripts) in the Interaction group of the Venn diagram for genes with log 2-fold changes greater than 1 for p-values ≤ 0.05 (pairwise pLSD) at each hourly time point. (e) Pathway overrepresentation analysis was performed with DAVID (Sherman et al., 2022) using Gene Ontology (GO) (Mi, Muruganujan, Ebert, Huang, & Thomas, 2019) analysis of the 1254 genes in the Interaction group. All data for the 5169 RNA seq transcripts that passed filtering metrics are shown in File S1 which also contains the two-way ANOVA results which can be used to sort for genes in each sector of the Venn diagram.

We analyzed the 1254 genes in the TxDxI group at each time point for statistically significant changes (criteria-p ≤ 0.05 Fisher’s protected Least Significant Difference pairwise contrast-pLSD, and fold change ≥ 2) (File S2). A notable result is that few genes are up-regulated until the 5 and 6 h time points and very few genes are down-regulated at any time point (Figure 1D). As we discuss in detail below, up-regulation of genes starting between the 4^th^ and 5^th^ h occurs when the majority of cells enter their second cell division cycle. Thus, these data explain much of the effect of time and drug components of the interaction gene set.

We recently reported that Myr treatment has a notable effect on the size of most amino acid pools which remain significantly smaller in drug-treated cells (N. L. Hepowit et al., 2021). To determine if transcription has roles in lowering amino acid pools and to gain a global view of significant features of the transcriptome, we performed a gene ontology (GO) analysis of genes in the TxDxI sector of the Venn diagram. GO terms Membrane and Plasma Membrane are highly enriched (Figure 1E), consistent with Myr slowing synthesis of sphingolipids along with synthesis of other lipids and membrane-bound proteins in the ER and consequent remodeling of membrane composition throughout the cell. Other high scoring terms include Actin Cortical Patch, indicating membrane growth and remodeling as well as cell growth and division. The term Oxidation-Reduction Process primarily identifies chemical reactions in metabolic pathways which are captured also by the KEGG pathway term Metabolic Pathways. Lastly, Endocytosis, another high ranking KEGG pathway, has functional links to the terms Membrane, Plasma Membrane and Actin Cortical Patch and to Mup1, the major methionine transporter, presented below.

Enriched GO terms in three other sectors of the Venn diagram (Figure 2a) were analyzed starting with genes listed in columns labeled Time x Drug and Time (Figure 2b). Our focus was on genes matching one of the 10 indicated patterns over the 1-6 h time-frame. Patterns were sorted into ones having transcripts that behaved like the diagramed pattern or like its mirror image (+ or –, respectively, as indicated in the column labeled ‘Corr’). Only patterns showing significantly more genes (indicated by an asterisk, binomial test ρ ≤ 0.05) assigned to them than expected by chance were analyzed. Most enriched GO terms in the Time x Drug group represent processes requiring many genes including cell growth and division (Figure 2c). The genes and GO terms present in all of the patterns analyzed are found in File S2. A graph of any gene transcript across the 1-6 h time-frame can be plotted using the Graph Reporter tab in File S1.

**FIGURE 2.**
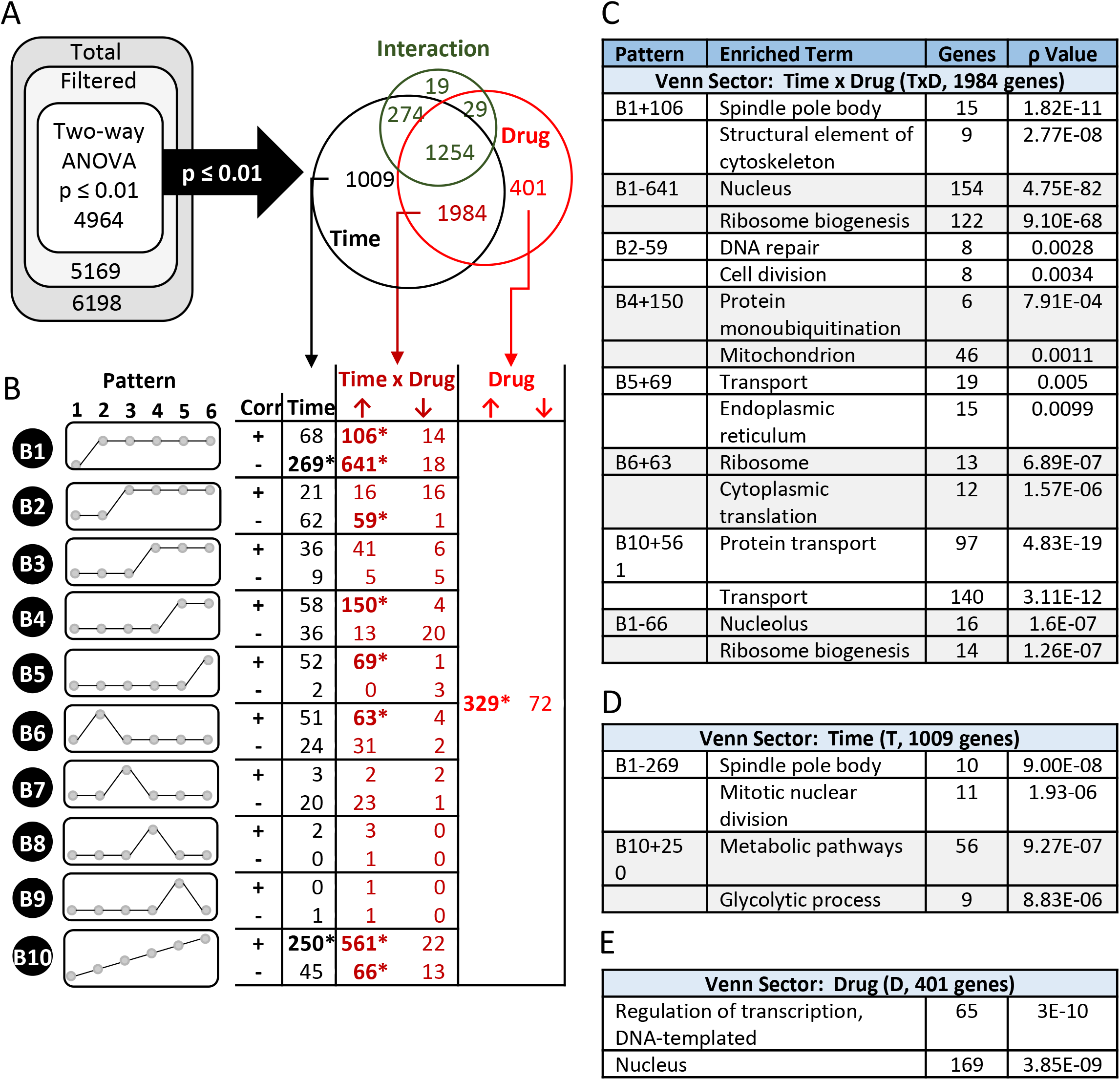
Detailed analysis of the Drug, Time and Drug x Time groups of the transcriptomics data set identified by two-way ANOVA analysis. (a) Diagram of two-way ANOVA analysis with results summarized in a Venn diagram showing the number of genes in each sector and indicating the three groups of genes (Time, Time x Drug and Drug) that were analyzed further for ones matching specific patterns. (b) Summary of genes matching the 10 indicated patterns. The column labelled “Corr” indicates genes having the same time course (+) or the inverse (mirror image) time course (-) of a pattern. Columns labeled Time, Time x Drug and Drug indicate the number of genes matching the corresponding pattern or its mirror image. Arrows at the top of the columns indicate genes up (↑) or down-regulated (↓) by drug. Statistically significant groups of genes are indicated by bold fond and with an asterisk (*). Genes and GO analysis of groups indicated by an asterisk are listed in File S2. (c - e) Summary of top enriched GO terms found in specific patterns.

In contrast to the Time x Drug sector, the 1009 genes in the Time sector of the Venn diagram fall into multiple patterns with the B1 and B10 patterns being most highly represented. Genes in this sector are not significantly changed by Myr but are significantly changed with time and serve as an example of time-related changes. All of these patterns will require further effort to determine their significance in Myr-induced longevity.

The Drug sector of the Venn diagram contains 401 genes with most (329, File S2) being up-regulated. As expected, these genes do not strongly associate with any temporal pattern (Figure 2b, right-most column). However, these genes do respond to Drug but not to Time. Because of our experimental design, we cannot be sure if some genes in this sector are driving the changes associated with time that we see but they could be. The most enriched GO terms are Regulation of Transcription, DNA Templated and Nucleus (Figure 2e), indicating a strong response to Myr treatment independent of time in culture.

In summary, GO term analysis of transcriptional changes in ten patterns during the 1-6 h time course of Myr treatment gives little indication of transcriptional involvement in keeping amino acid pools low in drug-treated cells. If there is a contribution, it may be overlooked because the number of genes involved is too small to achieve statistical significance at the pathway level, or transcriptional changes do not fall into any of the 10 patterns we examined.

### 2.2 Correlation analysis of transcriptomics data identifies clues to amino acid metabolism

As another approach to understand effects of Myr on the transcriptome, we analyzed transcript data using the R program package of Weighted Gene Correlation Network Analysis, WGCNA (Langfelder & Horvath, 2008), to identify clusters (modules) of highly correlated genes showing a similar response to Myr over the 1-6 h time course. A gene dendrogram produced by average linkage of hierarchical clustering revealed many modules with the seven most correlated having from 18 to 2490 genes (File S3, column L). Examination of the relationship of the modules to free amino acid pools over time identified the Brown and Green modules with 1126 and 144 genes, respectively, as having a negative correlation to Myr treatment: Myr maintains low pool levels but enhances the level of a transcript at one or more time points (Figure 3a). A network representing the relationship between genes in the Brown and Green modules includes the Turquoise module whose genes interact with both the Brown and Green modules (Figure 3b).

**FIGURE 3.**
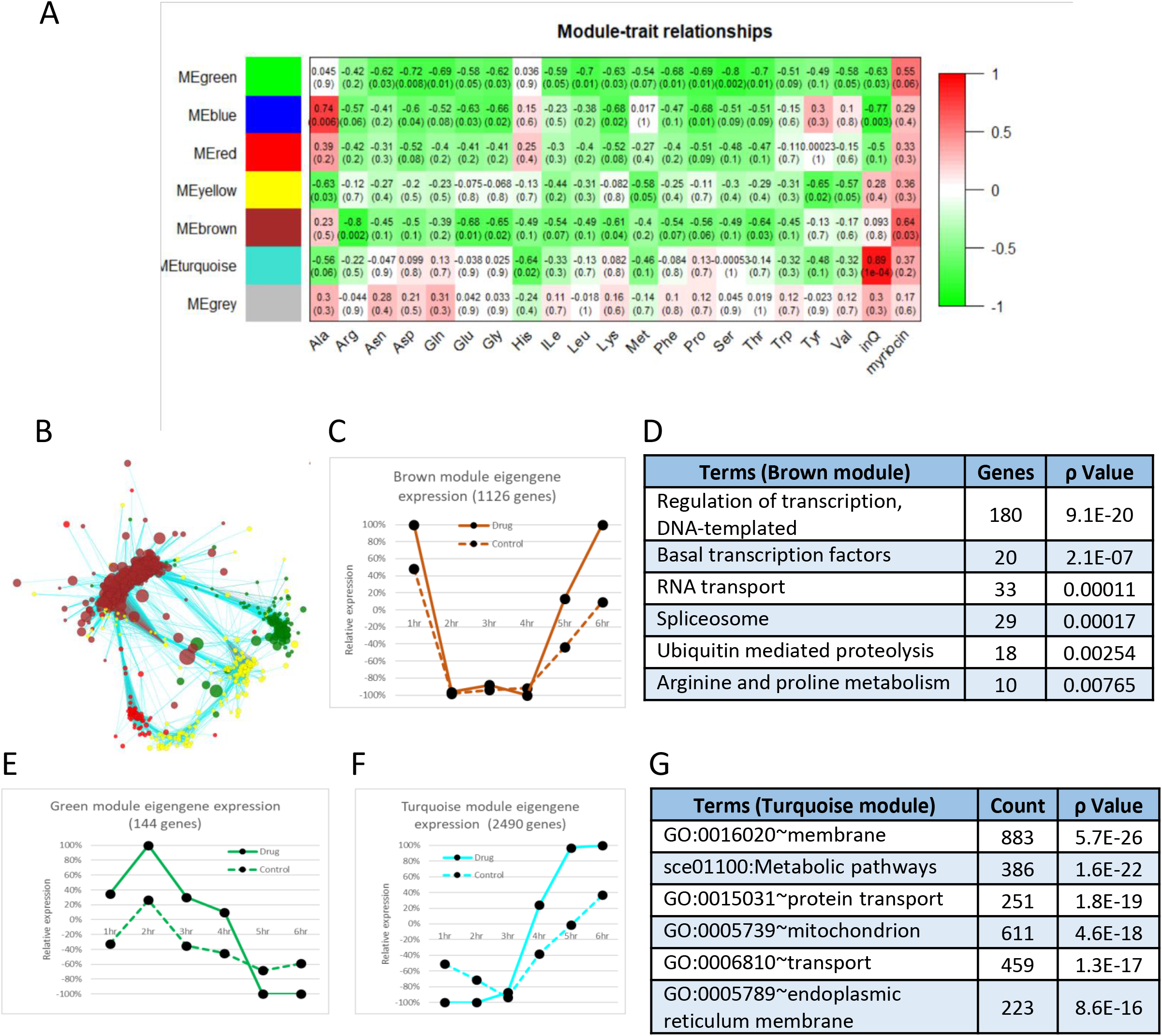
Correlation analysis of transcriptomics data. (a) The 7 color-coded modules whose gene members are highly correlated over time (left-hand side of diagram) were analyzed for their relationship to each amino acid pool (pool data are from (N. L. Hepowit et al., 2021)). The degree of correlation is indicated by the red-green (correlated-anticorrelated) scale at the right-side of the diagram. This figure indicates that the Green and Brown modules are the most anti-correlated (red shading) with amino acid pool, most of which are lowered by Myr treatment (right-most column labeled “myriocin” at the bottom of the diagram). Accordingly, genes in the Green module are up-regulate by myriocin treatment whereas pool size is down regulated. (b) Network diagram showing the relationship of genes in the Green and Brown modules which are connected by genes in the Turquoise. Genes are indicated by Nodes (circles) and relationships by edges. All genes and relationship values are presented in File S3. (c) Scatter plot of the Brown Eigengene across the 1-6 h time frame. (d) Enriched GO terms found in the Brown module. (e) Scatter plot of the Green Eigengene across the 1-6 h time. (f) Scatter plot of the Turquoise Eigengene across the 1-6 h time frame. (g) Enriched GO terms found in the Turquoise module. Genes used in calculating the mean Eigengene are in File S3 and the 7 mean Eigengene values and their scatter plots are presented in File S4.

Such negative correlation in a module is exemplified by an Eigengene representing the average effect of Myr on genes compared to the untreated drug control (File S4). For example, the Eigengene representing the Brown module shows that Myr induces transcription starting at 4 h (Figure 3c) whereas there is no increase in amino acid pool size in drug-treated cells (N. L. Hepowit et al., 2021). The highest-ranking GO term in the Brown module is Regulation of Transcription where drug treatment enhances transcript levels particularly after 4 h, consistent with the ANOVA analysis represented in Figure 1d. A different negative correlation is seen for the Eigengene representing the small Green module where genes are up-regulated by Myr from 1-4 h and then drop below values in untreated cells (Figure 3e). The main GO term in the Green module is Translation (Biological Process, p = 1.7E-88) along with Ribosome Assembly and related processes responding to drug-induced slowing of protein synthesis and growth rate (N. L. Hepowit et al., 2021). Lastly, the large Turquoise module with 2490 genes captures Myr-induced transcriptional events involving processes or pathways or cell components each with more than 200 genes (Figures 3e, g. Interestingly, transcripts represented in the Turquoise module rapidly increases after 4 h of drug treatment, substantiating the ANOVA analysis represented in Figure 1d. Substantial effort will be required to delineate the contribution of these genes to Myr-enhanced longevity.

### 2.3 Deubiquitination of Mup1 impairs Myr-enhanced longevity

A drawback of WGCNA and pathway and pattern analyses is a reduced ability to identify significant biological features involving smaller numbers of genes or for genes that do not fit a specific pattern across the 1-6 h time-frame. To circumvent these limitations and to discover transcript changes with potential roles in lowering amino acid pools, we examined genes with the highest possible significance (1E-16) in the Drug, Time and Interactions columns of data from the two-way ANOVA analysis of mRNA levels. This approach identified only 16 genes out of the 4964 genes (File S1, Filter Tab, columns M, N and O). The *UBI4* gene encoding the major stress-associated ubiquitin captured our attention because stresses of various types have known roles in aging and longevity and we previously identified stress responses in Myr-treated cells (Huang, Liu, & Dickson, 2012; Liu et al., 2013). A defining feature of *UBI4* transcript abundance in our studies is a rapid increase over the 4-6 h time period (Figure 4a, statistical significance of Area Under Curve (AUC) 95% CI (difference of the means): -189.081 to - 137.609, ρ-value = 0.00006). The path of the UBI4 transcript level corresponds to the time period in which Myr has its most significant effect on transcription (Figure 1d). The trajectory of *UBI4* transcripts appear to be like transcript pattern B4 with 150 genes up-regulated between 4-5 h in the Time x Drug column of Figure 2b. The highest-ranking GO term in this group of genes is Protein Monoubiquitination (Figure 2c): Protein Polyubiquitination is also found at a lower significance level in this group. However, due to the stringent nature of ANOVA analysis, the *UBI4* transcript does not fit in the B4+ pattern. But it does fits a unique pattern containing 72 genes (File S5). Importantly, the GO terms Protein Monoubiquitination and Protein Polyubiquitination are also enriched in this group. The potential biological significance of the *UBI4* transcript pattern may relate to our previous analyses of Mup1-pHluorin trafficking. We found that the fluorescent signal decreases in the PM more rapidly during the 4-7 h time-frame in Myr-treated cells, in a dose-dependent manner, which requires ubiquitin conjugation for endocytosis to occur (N. L. Hepowit et al., 2021).

**FIGURE 4.**
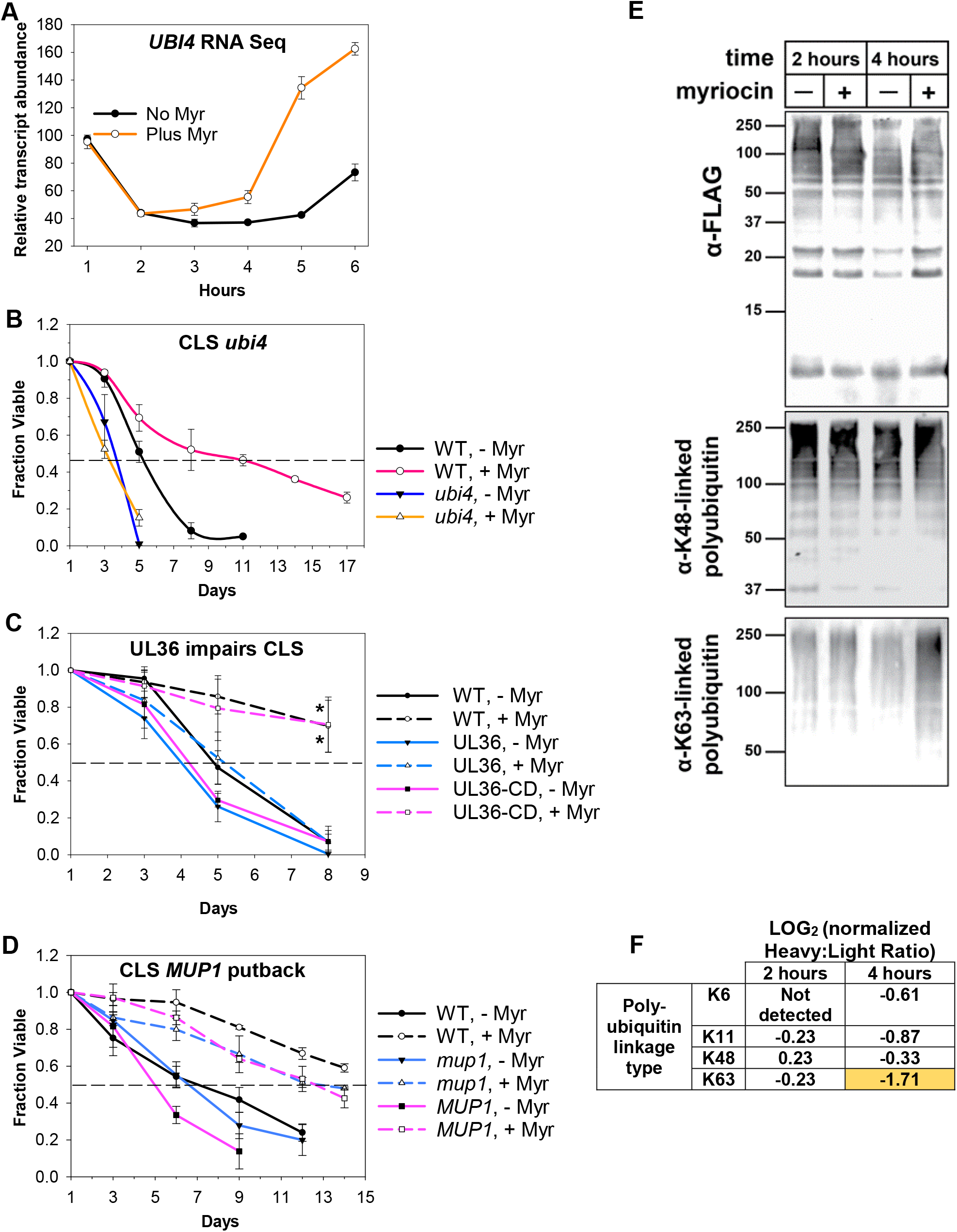
Ubiquitin plays a central role in Myr-enhanced longevity. (a) Summary of the relative abundance of *UBI4* transcripts across the 1-6 h time frame in Myr-treated or untreated cells. (b) CLS assay showing decreased survival of untreated (-Myr) *ubi4Δ* cells verses WT (BY4741) untreated cells and the complete lack of Myr-enhancement of lifespan in *ubi4Δ* cells compared to strong enhancement in WT cells. Error bars: SD (N = 3). (c) Data showing that removing ubiquitin from Mup1 impairs Myr-enhanced CLS. Cells having Mup1-pHlurion tagged with a UL36 deubiquitinase domain (DUB) are not able to respond to Myr treatment and enhance CLS. In contrast, replacing catalytically active UL36 with a catalytically dead UL36 domain (UL36-CD) restores Myr-enhanced lifespan. WT = BY4741 Mup1-pHluorin cells. Statistical significance at day 8 was determined (Student’s t-test) for Myr-treated WT vs UL36 and UL36-CD vs UL36 cells: ρ-values ≤ 0.024 and 0.032, respectively. Error bars: SEM (N = 3). (d) Lifespan assays reveal that deletion of *mup1* does not impair Myr-enhanced longevity. WT (BY4741) and *MUP1* putback (*mup1* allele replaced by *MUP1*) are control strains for the presence of a functional *MUP1* gens. Error bars: SEM (N = 3). (e) Affinity purified ubiquitin (from a yeast strain encoding N-terminally FLAG-tagged ubiquitin at two native, chromosomal ubiquitin genes) and blotted for total ubiquitin (top panel) captured as well as K63- and K48-linked polyubiquitin (middle and bottom panels, respectively) in cells treated on not treated with Myr after 2 or 4 h of cell growth. (f) SILAC-MS analysis of untreated (Heavy) or myriocin-treated (Light) yeast cells at the indicated time point. Heavy:Llight ratio quantifies relative abundance between the two samples. In this experiment, a negative LOG value indicates increased abundance in the myriocin-treated sample (and vice versa). All measurements were normalized to the Heavy:Light ratio for total ubiquitin (unmodified peptides), which did not significantly differ with Myr treatment at either 2 or 4 h post-myriocin treatment.

To determine if ubiquitination has roles in Myr-enhanced chronological lifespan (CLS), we examined untreated *ubi4Δ* cells and found a faster loss of viability compared to untreated BY4741 cells (Figure 4b, 50% survival day 3 vs day 5, respectively). Additionally, the CLS of *ubi4Δ* cells treated with Myr does not increase and remains the same as untreated cells. Untreated wild-type (WT, BY4741) cells show 50% survival at day 5-6 which is extended to day 11 by Myr treatment (AUC, 95% CI: -6.828 to -2.840, ρ-value = 0.0025). From these data we conclude that *UBI4* is required for Myr to enhance CLS.

Since ubiquitin has many functions, we studied its role in ubiquitin-mediated endocytosis of Mup1 using the same strains as used in our published analyses of endocytosis (N. L. Hepowit et al., 2021). Specifically, CLS was assayed using cells with chromosomal *MUP1* tagged with pHluorin-UL36 (catalytic active or catalytic dead, CD). UL36 is a viral deubiquitinase (DUB) that can reverse localized ubiquitin conjugation activity, thus protecting the fusion protein from ubiquitin-mediated degradation or trafficking events (MacDonald, Buchkovich, Stringer, Emr, & Piper, 2012). CLS is strongly enhanced by Myr treatment in two of these strains, namely WT cells (BY4741, *MUP1*-pHluorin) and cells with the catalytically dead UL36-CD domain (*MUP1*-pHluorin-UL36-CD) (Figure 4c, AUC of Myr-treated vs untreated cells, 95% CL: -2.710 to - 1.018, ρ-value = 0.0036 for WT and CI: -3.450 to -1.346, ρ-value = 0.0032 for UL36-CD). These significance values are probably an underestimate because both strains tend to regrow or gasp after day 8 which is why data are not shown beyond this time. In contrast, CLS is only slightly enhanced by Myr treatment in cells with a catalytically active UL36 domain (Figure 4c, AUC, 95% CI: -1.864 to -0.0284, ρ-value = 0.05). Statistical significance was also evaluated by using a t-test for the day 8 survival values for Myr treated cells: WT vs UL36 (ρ-value = 0.024) and UL36-CD vs UL36 (ρ-value = 0.032). Taken together, these data show that Myr-enhanced longevity depends on Mup1 ubiquitination.

To establish that deubiquitination is required for drug-enhanced longevity rather than the lack of Mup1 transport activity, we compared the CLS of *mup1Δ* cells to wild-type cells and cells with the *mup1Δ* allele replaced with *MUP1*. Myr treatment significantly enhanced CLS in each of the three strains (Figure 4d, for WT BY4741, *mup1Δ* and *MUP1* the AUC for Myr-treated vs untreated cells, ρ-values ≤ 0.009, 0.026 and 0.0024, respectively). The viability curves for untreated *mup1Δ* and *MUP1* cells do not differ significantly (AUC, 95% CI: -1.682 to 0.860, ρ-value = 0.42). We conclude from these data that deleting *MUP1* does not prevent Myr-enhanced longevity.

Given the increased *UBI4* transcript level after 4-6 h of Myr treatment (Figure 4a), we hypothesized that Myr treatment might affect total ubiquitin and it cellular distribution. To test this, we affinity purified FLAG-ubiquitin (from yeast cells harboring N-terminal FLAG fusions at two endogenous ubiquitin-encoding loci (*RPS31* and *RPL40B*)) from untreated or Myr-treated yeast cells. Immunoblot analysis revealed that Myr treatment for 2 h or 4 h did not significantly alter the amount of FLAG-ubiquitin recovered or the amount of K48-linked ubiquitin polymers recovered (Figure 4e). In contrast, we observed a significant increase in K63-linked ubiquitin polymers after 4 h of Myr treatment (Figure 4e, right lane of bottom immunoblot), suggesting that increased conjugation of ubiquitin in K63-linked polymers is part of the cellular response to depletion of sphingolipids.

To confirm this result, and to resolve other linkage types, we performed *<u>s</u>*table *i*sotope *l*abeling with *<u>a</u>*mino acids in cell *<u>c</u>*ulture (SILAC) on yeast cells and subjected light-labelled and heavy-labelled cells to mock-treatment and Myr-treatment, respectively. At 2 h and 4 h of treatment, we collected cells, prepared lysates, and affinity purified FLAG-ubiquitin, followed by mixing, tryptic digestion, and processing of peptides for analysis by mass spectrometry. This SILAC-MS analysis resolved various peptides corresponding to the different linkage types of ubiquitin polymers, including K6-linked, K11-linked, K48-linked, and K63-linked polymers. Interestingly, we detected only modest changes in linkage types after 2 h of Myr treatment, while 4 h of Myr treatment resulted in increased formation of several polymer types, most notably for K63-linked ubiquitin polymers (Figure 4f). This finding is consistent with our immunoblot results shown in Figure 4e, confirming that yeast cells increase the formation of K63-linked ubiquitin polymers in response to Myr treatment. Taken together, these results suggest that yeast cells respond to Myr treatment by remodeling ubiquitin pools, partly by increasing production of ubiquitin via transcription of *UBI4* and partly by deploying existing ubiquitin pools to promote increased formation of K63-linked ubiquitin conjugates.

## 3 DISCUSSION

We performed transcriptomics analysis for several reasons including (i) to determine if transcription played roles in keeping amino acid pools at a low level in Myr-treated cells, (ii) to determine if Myr treatment caused a major transcriptional shift during a specific time-frame and (iii) to search for novel processes or pathways required for Myr-enhanced longevity. From our analysis, we are able to draw several conclusions. First, Myr influences transcription of very few genes during the first 4 h of treatment when 50-60% of cells are acting in synchrony. Second, transcription is robustly up-regulated beginning around the 4th h of Myr treatment when the majority of cells advance through the second cell division cycle. Third, transcriptional changes are not the major force promoting initial amino acid pool lowering. Although we cannot altogether exclude a role for transcription, we showed previously (N. L. Hepowit et al., 2021) that amino acid pool lowering is at least partly due to inactivation of amino acid transporters at the PM following Myr treatment. Fourth, ubiquitination of the methionine transporter Mup1 at the PM is vital for Myr-enhanced longevity. Lastly, Myr treatment strongly up-regulates K63-linked ubiquitination of proteins. Importantly, we have also recently reported that reducing sphingolipid synthesis inhibits the endocytosis of many nutrient transporters, while specifically promoting endocytic clearance of the methionine transporter Mup1 (N. L. Hepowit, Moon, B. J., Dickson, R. C. & MacGurn, J. A., 2022). Taken together, our analysis reveals a complex cellular response to Myr that involves modulation of several biological processes including endocytic trafficking, proteostasis, and amino acid homeostasis.

We performed transcriptomics analysis for several reasons including (i) to determine if transcription played roles in keeping amino acid pools at a low level in Myr-treated cells, (ii) to determine if Myr treatment caused a major transcriptional shift during a specific time-frame and (iii) to search for novel processes or pathways required for Myr-enhanced longevity. Our previous work established that amino acid pools are lower in Myr treated cells compared to untreated cells after 1 h of treatment (N. L. Hepowit et al., 2021). In the studies reported here we do not identify any indication of initial amino acid pool lowering being driven by transcription. However, we cannot eliminate possible effects on some amino acid pools. We were able to determine when cells respond transcriptionally to drug treatment under our assay conditions (Figure 1a). Transcripts (genes) changing in a statistically significant way across the initial 6 h of Myr treatment were identified by two-way ANOVA analysis (Figure 1b). Significant genes were sorted into ones responding primarily to Time (T) or Drug (D) or Interaction (I) (Figure 1c). To generate an overview of drug effects over time, we examined the 1254 genes in the TxDxI group at each time point for genes with 2-fold changes greater or less than 1 (Figure 1d). Surprisingly, very few transcripts that are significant by Interaction respond to Myr until after 4 h of treatment which, for the majority of cells, occurs during the second cell division cycle. The major GO terms in the TxDxI sector of the Venn diagram reflects processes, reactions and compartments related to cell growth and division (Figure 1e). The large upsurge of transcripts after 4 h of Myr-treatment along with the cellular processes and compartments they represent was also identified by correlation analysis as visualized by the Eigengene for the Brown and Turquoise modules (Figures 3c, f). The Turquoise module being especially significant as it contains nearly half of all transcripts identified as significant by two-way ANOVA analysis (Figure 1a). Our data probably underestimate the number of significant genes at each time point since only 50-60% of cells are acting in synchrony (N. L. Hepowit et al., 2021) or that temporal blurring distorted temporal assignment. Nonetheless, these possibilities are not likely to change our conclusion that Myr causes its greatest effects on transcription after about 4 h of treatment or after cells have executed their first cell division.

Possible future studies to better understand Myr-effects on amino acid and other types on metabolism are implicated by our transcriptomics data. The GO term Metabolic Pathways (193 genes) was identified in the Interaction analysis as being highly significant across the 1-6 h time-frame (Figures 1d, e). GO term analysis of these 193 genes identified KEGG pathways for carbon metabolism, secondary metabolites, amino acid metabolism (tryptophan, methionine, lysine, arginine, proline, histidine) and other types of metabolism as being significantly enriched (File S2, Tab “TxDxI 1254”). Several of these pathways have known roles in longevity (e.g. glycogen and trehalose metabolism), suggesting that the Myr-sensitive pathways defined in this work are novel but include elements of pathways defined in prior work as having roles in longevity. Endocytosis is another GO term found to be highly enriched in the Interaction analysis of transcripts across the 1-6 h time-frame (Figures 1d, e). GO term analysis of this group of 39 genes shows enrichment for genes involved in ubiquitin-mediated endocytosis which provide clues for the proteins, lipids and cellular machinery controlling Myr-induced endocytosis of Mup1 that we previously observed (N. L. Hepowit et al., 2021). The 39 genes in this group will facilitate understanding the cellular processes controlled by Myr that depend on ubiquitin and deubiquitination of Mup1 and that contribute to Myr-enhanced lifespan (Figures 4b, c). Finally, while Myr treatment promotes lowering of amino acid pools by reducing amino acid uptake (N. L. Hepowit et al., 2021), additional mechanisms may be vital to maintain amino acid pools at a level that balances a reduced rate of cell growth - which occurs at around 4 hr of drug treatment – with cellular remodeling necessary to transition to a quiescent state. One such mechanism may involve specific types of autophagy since transcripts in this processes are significantly represented in the Interaction component (1254 transcripts) of our transcriptomics data (Venn diagram, Figure 1c, d). Up-regulating autophagy could not only supply amino acids in drug-treated cells, but it could also foster proteostasis and removal of damaged organelles, a hallmark of longevity (Lopez-Otin, Blasco, Partridge, Serrano, & Kroemer, 2013).

Healthy proteostasis relies on proper functioning of protein degradation networks to maintain proteome quality control and prevent accumulation of misfolded and damaged proteins. Thus, a decline in proteostasis is a hallmark of aging and age-related diseases (Lopez-Otin et al., 2013). Analysis of the yeast transcriptional response to Myr treatment revealed a significant induction of the mRNA transcript encoded by *UBI4* (Figure 4a) – one of four genes in the yeast genome encoding for ubiquitin. The other three (Finley, Ozkaynak, & Varshavsky, 1987) ubiquitin genes (*RPS31, RPL40A*, and *RPL40B*) encode ubiquitin fusions to ribosomal subunits and generate most cellular ubiquitin in normal growth conditions, which inherently couples the translational and degradative capacities in the cell. By comparison, *UBI4* encodes a linear (heat-to-tail) ubiquitin pentamer that does not contribute significantly to the total pool of ubiquitin in normal growth conditions. However, in conditions of heat stress, oxidative stress, or starvation *UBI4* is transcriptionally induced which serves to increase the degradation capacity of the cell and uncouple it from ribosome biogenesis and translational capacity (Cheng, Watt, & Piper, 1994; Finley et al., 1987). Thus, our observation that Myr induces transcription of *UBI4* but not the other genes encoding ubiquitin (transcript levels of other ubiquitin genes can be displayed by using the ‘Reporter’ tab in File S1) is suggestive of a proteotoxic and/or starvation stress response involving activation of protein degradation networks and sizeable proteome remodeling. The importance of *UBI4* in Myr-enhanced longevity is underscored by our finding that the CLS of untreated *ubi4Δ* yeast cells decreased compared to WT cells and was not significantly enhanced by Myr treatment (Figure 4b). In addition, our finding that cells with chromosomal *MUP1* tagged with pHluorin-UL36 fail to show Myr-enhanced CLS (Figure 4c) suggests a novel role for ubiquitination in longevity. Importantly, we previously reported that UL36 fusion to Mup1 impedes its Myr-induced endocytosis (N. L. Hepowit et al., 2021), underscoring the linkage of ubiquitin-mediated endocytosis of Mup1 to Myr-mediated longevity. One limitation of this analysis is the potential for proximal effects on other proteins in the vicinity of the Mup1-pHluorin-UL36 protein (MacDonald et al., 2012; Stringer & Piper, 2011). Thus, we cannot exclude the possibility that ubiquitination of Mup1-associated factors is critical for Myr-mediated longevity. Future studies will be required to evaluate if such proximal effects are involved in the impairment of Myr-enhanced lifespan. The outcomes of these studies will potentially provide new targets or strategies for improving human healthspan.

## 4 METHODS AND MATERIALS

### 4.1 Strains, culture conditions, lifespan assays and statistical significance

Strains (Table 1), culture conditions, lifespan assays and their statistical significance were similar to ones described previously (N. L. Hepowit et al., 2021). CLS assays were performed at least twice using triplicate cultures. Concentrations of Myr used in CLS assays ranged from 475-525 ng/ml depending on the sensitive of strains compared to wild-type, prototrophic BY4741. Drug-sensitivity was measured by culture density (A600nm) after 24 h of growth using cultures started at 0.15 A600nm units of cells/ml. All BY4741 yeast strains used for lifespan assays and for RNA extraction were made prototrophic by transformation with the pHLUM plasmid (Addgene, Watertown, Massachusetts) (Mulleder et al., 2012). The DUB (UL36) fusion yeast strains used in this study were generated by homologous recombination in BY4741 and SEY6210 background strains using the reagents and strategy previously described (Hepowit et al. 2022). The *MUP1* knock-in strain (NHY945) was generated in BY4741 by swapping the endogenous *MUP1* coding region with NATMX (NHY930), shuffling the chromosomally integrated *NATMX* with *URA3* (NHY938.1), and snipping out *URA3* by homologous reintegration of a PCR-amplified *MUP1* coupled with counter selection on 5-fluoroorotic acid (5-FOA) synthetic media plate. A two-tailed Student’s t-test and Area Under the Curve (AUC)(Sigma Plot) were used to evaluate statistical significance of CLS assays done with at least 3 biological replicates. Results were verified by one or more repeat experiments.

**Table 1.**
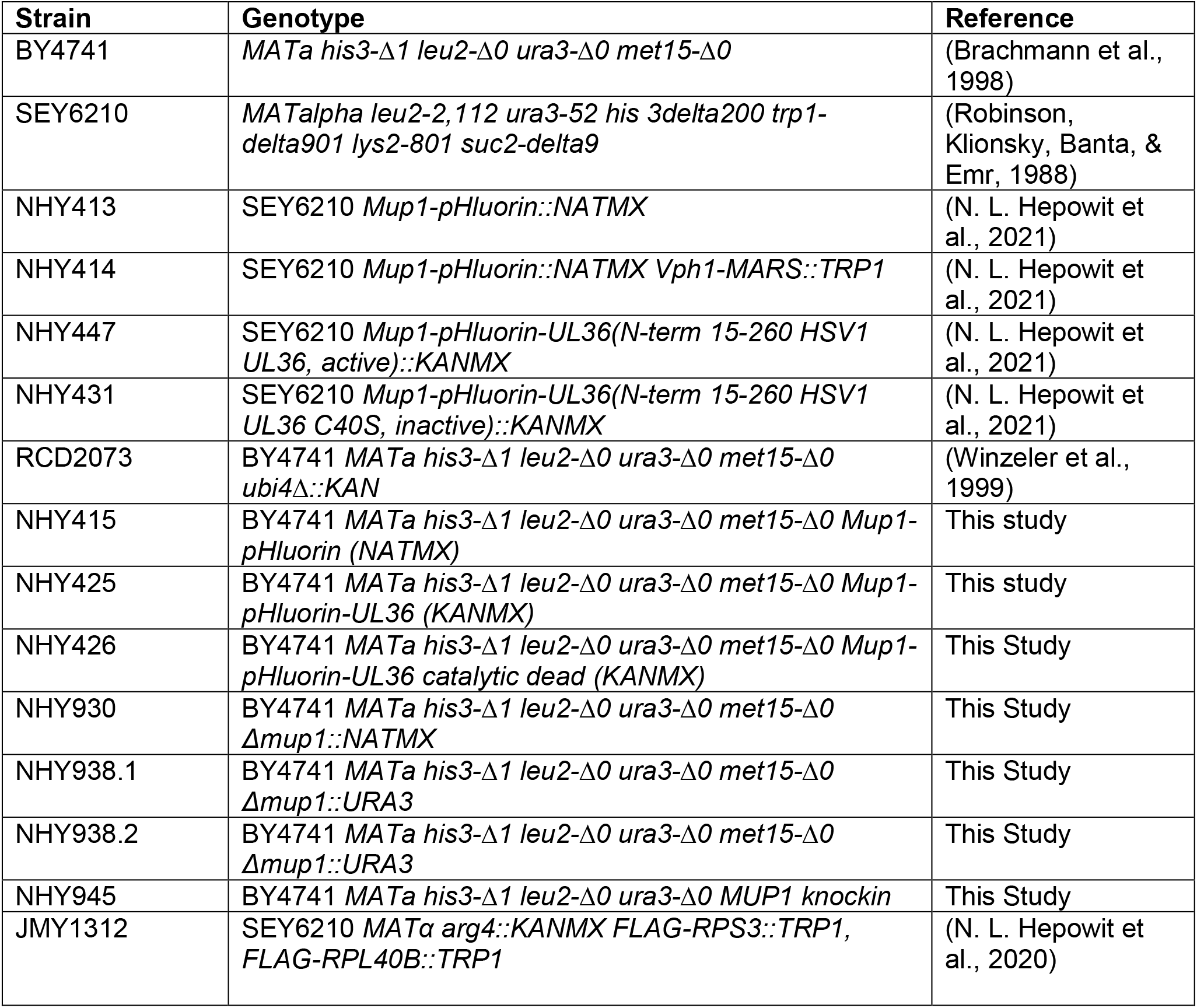
Strains used in this study.

Budding yeast mRNA-enriched profiles were generated from 42 individual samples of prototrophic BY4741 cells, including 3 replicates at 7 time points and 2 treatments (control and drug-treated). Culture conditions were similar to ones described previously (Hepowit et al. 2021, Aging) except for the following modifications. Prototrophic BY4741 yeast cells were grown in 200 ml of SDC culture medium in a 1 L flask with the medium heated to 30°C before addition of EtOH (final concentration of 0.3% for control samples). For drug-treated samples, myriocin was added after addition of EtOH to give final concentrations of 0.3 % EtOH and 700ng/ml myriocin. Lastly, cells from a saturated overnight culture were diluted into the 200 ml cultures to give an initial A600nm units/ml of 0.15. Flasks were incubated at 30°C and 200 rpm on a rotary shaker. Control and myriocin-treated cells were harvested at time 0, 1, 2, 3, 4, 5 and 6 h (all time points for each replication were from the same culture flask).

### 4.2 RNA extraction

RNA was extracted from 5 A600nm unit/ml of yeast cells harvested by filtration on a membrane filter at each time point as described previously for amino acid extraction (N. L. Hepowit et al., 2021). Filtered cells were washed once with 5 ml of ice-cold nanopure water and the filter was quickly transferred to a chilled 1.5 ml microfuge tube containing 0.5 ml cold nanopure water. Tubes were vortexed 10 sec followed by centrifugation for 15 sec. Supernatant fluid (450 μl) was transferred to a new tube and frozen in a dry-ice EtOH bath followed by storage at -80°C. Cold acid-washed glass beads (300 μl, 0.5 mm dia.) were added to a frozen cell pellet followed by addition of 300 μl of RLT buffer (RNAeasy mini kit, Qiagen, Germantown, Maryland). Tubes were vortexed 5 min at room temperature and placed on ice for 1 min. After 4 cycles 300 μl of ice-cold RLT buffer was added followed by mixing and centrifugation at 13,000xg for 2 min at room temperature. Supernatant fluid (450 μl) was transferred to a new microfuge tube and then mixed with 1 ml of 70% EtOH before transfer to a RNeasy spin column and processed according to the manufacturer’s instructions. Processed samples were frozen in a dry-ice EtOH bath and stored at -80°C. RNA seq was performed on total RNA samples at the Roy J. Carver Biotechnology Center at the University of Illinois.

### 4.3 RNA seq analysis

RNA sequencing was performed on total RNA samples at the Roy J. Carver Biotechnology Center at the University of Illinois. Two different mixes of ERRC spike-in RNA controls were added to the samples; one mix for the control samples, and one mix for the drug-treated samples. Libraries were constructed with Illumina’s ‘TruSeq Stranded mRNAseq Sample Prep kit’. Each library was quantitated by qPCR and sequenced on one lane for 151 cycles from each end of the fragments on a HiSeq 4000 using a HiSeq 4000 sequencing kit version 1. Fastq files were generated and demultiplexed with the bcl2fastq v2.17.1.14 Conversion Software (Illumina). Each sample’s pair of fastq files were run through trimmomatic 0.36 to first remove any remaining standard Illumina PE v3 adapters, then trim bases from both ends with quality scores below 28, and finally to remove individual reads shorter than 30 bp and their paired read, regardless of length (parameters ILLUMINACLIP:/home/apps/trimmomatic/trimmomatic-0.36/adapters/TruSeq3-PE.fa:2:15:10 TRAILING:28 LEADING:28 MINLEN:30). Paired reads per sample were pseudo-aligned to the Yeast R64 transcriptome and 92 ERRC spike-in control sequences using Salmon 0.8.2 with parameters -l A --numbootstraps=30 --seqBias –gcBias. Resulting FASTQ files mapped to the yeast genome (R64), resulted in count files and normalized using the transcripts per million (TPM) algorithm (Trapnell et al., 2010) using WebMev (Wang, Kutnetsov, Partensky, Farid, & Quackenbush, 2017). Resulting data were downloaded as flat files and loaded into Excel for further analysis. From a total of 6198 mapped genes, 5169 were uniquely annotated with gene symbols and had sufficient non-zero readings for further analysis. The filtered data were analyzed by two-way ANOVA for the main effects of drug and time, as well as for interaction. Significant by the time term or both the drug and time term, data were further analyzed by post-hoc pairwise Fisher’s protected Least Significant Difference (pLSD), and log 2-fold change comparison to further isolate the effects of drug over time. Template analysis was applied using pre-determined templates (depicted in Figure 2) correlated to average expression over time in drug or control conditions. Each gene significant by time and/or interaction was correlated to each of the ten templates and was assigned to the template with which it had the strongest correlation. Data have been deposited in the GEO (GSE199904) [NCBI tracking system #22817261]. GO terms were identified by using DAVID software (Sherman et al., 2022).

### 4.4 Gene regulatory network using WGCNA

The WGCNA (v1.70-3) (Langfelder & Horvath, 2008) was used to identify gene modules and build unsigned co-expression networks, which include negative and positive correlations. Briefly, WGCNA constructs a gene co-expression matrix, uses hierarchical clustering in combination with the Pearson correlation coefficient to cluster genes into groups of closely co-expressed genes termed modules, and then uses singular value decomposition (SVD) values as module eigengenes to determine the similarity between gene modules or to calculate association with sample traits (for example, incubation time or amino acid levels). The top 2,000 variable genes were used to identify gene modules and network construction. Soft power 8 was chosen by the WGCNA function pick SoftThreshold. Then, the function TOMsimilarityFromExpr was used to calculate the TOM similarity matrix via setting power = 8, networkType = “signed. The distance matrix was generated by subtracting the values from similarity adjacency matrix by one. The function flashClust (v.1.01) was used to cluster genes based on the distance matrix, and the function cutreeDynamic was utilized to identify gene modules by setting deepSplit =3. Cytoscape (v.3.8.2) was applied for the network visualizations.

### 4.5 Analysis of ubiquitin and its linkage types

For immunoblotting assays 5 A_600_ unit of cells were precipitated in 10% trichloroacetic acid on ice for 30 min. The protein precipitate was washed twice with ice-chilled acetone, lyophilized by vacuum centrifugation, and resuspended in urea sample buffer (75 mM Tris-HCl [pH 6.8], 6 M urea, 1 mM EDTA, 3% SDS, 20% glycerol, bromophenol blue), heated at 65°C for 5 min, and vortexed for 5 min. Proteins were separated by SDS-PAGE and transferred onto Immobilon-P^SQ^ membrane (0.2 µm; Millipore). Immunoblotting was performed using the following primary antibodies: anti-ubiquitin (1:10,000; LifeSensors; MAb; clone VU-1), anti-K48 (1:10,000; Cell Signaling; RAb; clone D9D5), and anti-K63 (1:4000; EMD Millipore; RAb; clone apu3). Secondary antibodies were anti-mouse (IRDye 680RD-Goat anti-mouse) or anti-rabbit (IRDye 800CW-Goat anti-rabbit) obtained from LI-COR Biosciences). Blots were scanned using Odyssey CLx and signal fluorescence was visualized using Image Studio Lite (LI-COR Biosciences).

Ubiquitin linkage types in JMY1312 yeast cells were determined and quantified by SILAC-based mass spectrometry as previously described except for a change in the cell lysis buffer: 50 mM Tris-HCl [pH7.5], 150 mM NaCl, sodium pyrophosphate, 20 mM β-glycerophosphate, 2 mM sodium orthovanadate, 1 mM phenylmethylsulfonyl fluoride, 0.2% NP-40, 10 mM iodoacetamide, 20 µM MG132, 1 mM 1,10-phenanthroline, 1X EDTA-free protease inhibitor cocktail (Roche), 1X PhosStop (Roche) (N. L. Hepowit, Pereira, Tumolo, Chazin, & MacGurn, 2020).

## Supporting information

Supplementary File 1_RNA seq.dataset

Supplementary File 2_Venn Diagram & pattern group GO analyses

Supplementary File 3_Cluster analysis

Supplementary File 4_Mean Eigengene graphs

Supplementary File 5_UBI4 group

## ACKNOWLEDGEMENTS

Research reported herein was supported by the National Institute On Aging of the National Institutes of Health under Award Number R56AG024377 (RCD) and the National Institute of General Medical Sciences of the National Institutes of Health under Award Number R35GM144112 (JAM). The content is solely the responsibility of the authors and does not necessarily represent the official views of the National Institutes of Health

## CONFLICTS OF INTEREST

The authors declare that they have no conflicts of interest

## AUTHOR’S CONTRIBUTIONS

NLH, RCD and JAM designed the experiments. NLH and RCD conducted the experiment. EB performed the RNA seq analysis. SL performed the transcript correlation analysis. All authors were involved in data analysis. RCD and JAM wrote the manuscript.

## SUPPORTING INFORMATION

Additional supporting information may be found online

## Notes

### Competing Interest Statement

The authors have declared no competing interest.

### Summary of Updates

Revise Abstract and addition of more data and discussion of it.

